# Lipopolysaccharide and Wzi retain capsular polysaccharide at the *Klebsiella pneumoniae* cell surface and promote bloodstream infection fitness

**DOI:** 10.64898/2026.10.30.762281

**Authors:** Saroj Khadka, Zachary J. Resko, Grace E. Shepard, Emily L. Kinney, Gabriella M. Gates, Drew J. Stark, Laura A. Mike

## Abstract

Bacterial capsules are formed by individual capsular polysaccharide (CPS) polymers and are the outermost extracellular layers of many Gram-positive and Gram-negative bacteria. Capsules affect several aspects of bacterial biology, including resistance to host immune factors, survival in unfavorable environments and antimicrobial resistance. Research on CPS biology has largely focused on its synthesis and the pathogenic consequences of its presence or absence. Beyond synthesis, our understanding of its retention (attachment) on the cell surface is limited. *Klebsiella pneumoniae* is an important Gram-negative pathogen and expresses more than 80 capsular serotypes using the common Wzy-dependent CPS biosynthesis system and phenotypically manifests in various forms, making this species an attractive bacterial model to study the underexplored aspects of CPS biology. Here, we identified and characterized bacterial components involved in CPS retention in *K. pneumoniae* and assessed the importance of CPS retention on bloodstream fitness. Our examination revealed that lipopolysaccharide (LPS) and the outer-membrane protein, Wzi, retain *K. pneumoniae* CPS on the bacterial surface. We demonstrate that the core-oligosaccharide domain of LPS and the extracellular surface of Wzi likely provide anchoring sites for CPS on the bacterial surface. Furthermore, CPS retention depends on divalent cation availability. Importantly, CPS retention on the bacterial surface is important for survival in serum and bacterial fitness at secondary infection sites in a murine bacteremia model. Our findings altogether demonstrate that the functional importance of CPS to bacteria extends beyond synthesis and its overall abundance. Bacterial control over other CPS properties, like cell surface retention, are also important for *K. pneumoniae* fitness during bacteremia.

## 1. INTRODUCTION

*Klebsiella pneumoniae* is a leading cause of global morbidity and mortality, and attributed to nearly 800,000 deaths annually [1]. Due to rising antimicrobial resistance and the public health burden, drug-resistant *K. pneumoniae* has been classified as a high-priority pathogen requiring new therapeutic targets by the World Health Organization. Capsular polysaccharide (CPS) production is a near universal feature and a key virulence determinant across the members of *K. pneumoniae* species complex [2]. The role of CPS in preventing phagocytosis, limiting complement-mediated killing and promoting immune evasion is well-established [2]. However, CPS production and/or retention are not always advantageous to bacteria. First, CPS shedding can be an active strategy to misdirect host immune components as observed in *Streptococcus pneumoniae* [3]. In *S. pneumoniae*, bacterial autolysin LytA facilitates removal of CPS from the surface in response to cationic antimicrobial peptides (CAMP), and promotes invasion of lung epithelial cells [3]. A similar observation in response to CAMP have been reported for *K. pneumoniae* and *P. aeruginosa* [4]. Second, in *K. pneumoniae* specifically, persistently colonizing bacterial populations frequently accumulate spontaneous mutations that disrupt CPS synthesis, suggesting that there are niche-dependent evolutionary pressures selecting for CPS loss [5]. Third, CPS is a major surface recognition factor for bacteriophages, and as such CPS loss is a bacterial countermeasure to protect against phage infection. For example, in *K. pneumoniae*, CPS abundance is reversibly downregulated in response to phage infection [6]. Together, these findings make CPS and other surface polysaccharides attractive therapeutic targets and highlight the importance of understanding the underlying mechanisms of synthesis, export and anchoring [7,8].

*Klebsiella* species synthesize the high-molecular weight CPS polymers via the Wzy-dependent CPS biosynthesis system, which are exported out of the system and organized on the cell surface. While a substantial body of literature has described the synthesis and export process in detail, how these polymers are retained and organized on the cell surface remains incompletely understood. CPS retention has been proposed to involve dedicated CPS-binding protein Wzi or interactions with another abundant cell-surface polysaccharide lipopolysaccharide (LPS) [9,10]. In addition, strains expressing mutations in the CPS biosynthesis system components like Wzc, which functions as a regulator of CPS polymerization rate, synthesize increased proportion of exopolysaccharides that are not associated to the bacterial cell surface [11].

Based on literature, we investigated the bacterial factors determining *K. pneumoniae* CPS retention and the consequences of altered CPS retention on pathogenesis. We demonstrate that CPS retention is not dictated by a single dominant factor and instead arises from combined contributions of the components of CPS biosynthesis machinery, additional surface polysaccharides, dedicated CPS-binding protein and availability of divalent cations. Finally, we show that both CPS retention and LPS are critical fitness determinants during bloodstream infection, each distinctly contributing to resistance against serum factors and to persistence in blood-filtering organs.

## 2. RESULTS

### Capsular polysaccharide (CPS) retention in *Klebsiella pneumoniae* requires both core-lipopolysaccharide (core-LPS) and Wzi

LPS components and the cell-surface lectin Wzi have previously been implicated in *K. pneumoniae* CPS retention [10,12]. In our previous work, we additionally identified a spontaneous point mutation in the CPS chain length regulator Wzc that impairs CPS retention [11]. Building on this knowledge, we sought to validate and further define the roles of these cellular components in CPS retention using the hypervirulent model strain KPPR1 and its corresponding isogenic mutants. KPPR1 is a rifampin-resistant derivative of *K. pneumoniae* ATCC 43816 that synthesizes O1-type LPS and K2-type capsule, as confirmed by PathogenWatch (Kaptive v.3.1) [13,14].

Gram-negative LPS consists of three components: a variable O-antigen polysaccharide (O-PS) chain, a core polysaccharide (core-OS) chain comprising outer and inner regions, and the lipid A anchor that retains the LPS molecule in the bacterial outer membrane **(Fig 1A)** [15]. To examine the role of each LPS component in CPS retention, we generated KPPR1 mutants with sequential LPS truncations by deleting the ligase genes responsible for ligating the respective LPS components **(Fig 1A)**. Specifically, Δ*waaL* removed the O-PS from LPS, Δ*wabG* truncated both O-PS and outer core-OS, and Δ*waaC* truncated both O-PS and entire core-OS, leaving only the lipid A moiety **(Fig 1B)**. In contrast, the Δ*wzi* or the *wzc* point mutant (Wzc_Q395K)_ had a similar LPS profile as WT **(Fig 1B)**. Furthermore, uronic acid quantification revealed only a modest reduction in total CPS synthesis across all mutants, which reached statistical significance for Δ*wzi* **(Fig 1C)**.

**Fig 1:**
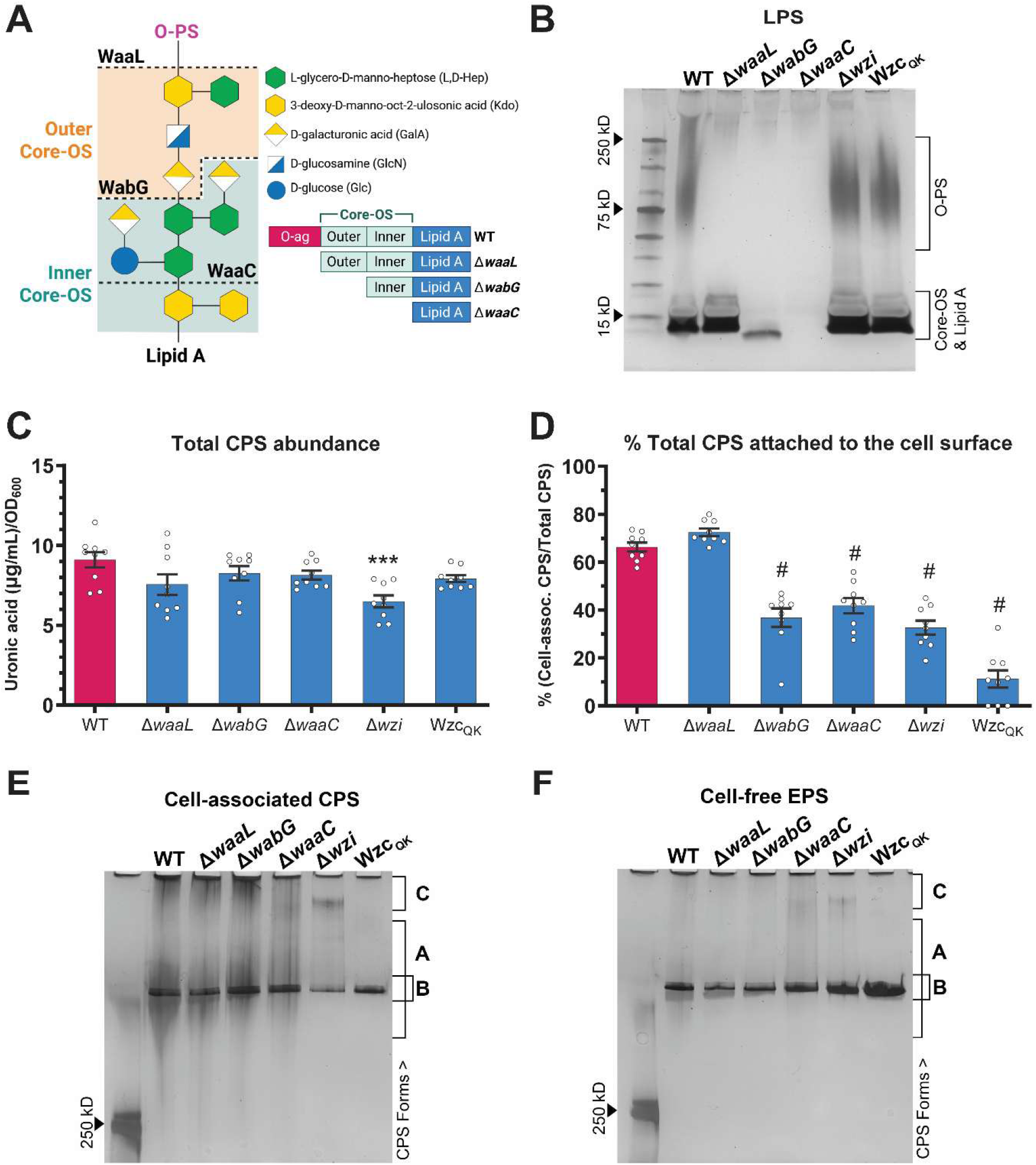
*K. pneumoniae* core-LPS and the cell-surface lectin, Wzi, are required for CPS retention on the outer membrane. *K. pneumoniae* strain KPPR1 wildtype (WT) and mutants were cultured in LB and their LPS and CPS were isolated and quantified by biochemical assay or visualized by staining the resolved polysaccharides via electrophoresis. KPPR1 mutants included isogenic deletions of LPS ligase genes (O-antigen ligase, *waaL*; outer-core ligase, *wabG* and inner-core ligase, *waaC*), deletion of cell-surface lectin gene *wzi,* and an isogenic mutant with single substitution in *wzc* residue Q395 (Wzc_QK)_. **(A)** Schematic of KPPR1 O1-type LPS with type-2 core showing the three principal LPS components: O-antigen (O-polysaccharide; O-PS), outer and inner core oligosaccharide (core-OS) and lipid A. ‘WaaL’, ‘WabG’ and ‘WaaC’ marked with dashed lines indicate the site of ligation catalyzed by the respective enzyme. The bottom-right illustration depicts the expected LPS structure of the LPS truncation mutants. The LPS structure was inferred from Whitfield et al [41]**. (B)** Crude LPS was resolved by SDS-PAGE and stained using silver stain. **(C)** Total CPS abundance was determined by measuring uronic acid content in total crude CPS extracts and normalized to OD_600._ **(D)** The percentage of total CPS retained on the cell surface was calculated as the ratio of cell-associated CPS to total CPS measured in (C). Exact cell-associated CPS and cell-free EPS uronic acid levels are available in S1B Fig. **(E)** Cell-associated CPS and **(F)** cell-free EPS were isolated, resolved by SDS-PAGE and stained with alcian blue followed by silver stain. ‘A’, ‘B’ and ‘C’ represent different CPS forms, categorized by their chain length, where ‘A’ refers to CPS chains with diverse length, ‘B’ refers to chains with uniform length and ‘C’ refers to ultra-high molecular weight of very-long CPS chains. Data presented are (B,E,F) the representative of three gel images or (C–D) mean of independent experiments with error bars representing the standard error of the mean. (C–D) Statistical significance was determined by comparing mutants to WT using one-way ANOVA with Dunnett’s post-hoc test. *P*-values are displayed above each bar; *** indicates *p* ≤ 0.001, # indicates *p* ≤ 0.0001. Experiments were performed three independent times, each with (B,E,F) single biological replicate or (C–D) biological triplicate (*n* = 9).

Among the LPS mutants, loss of the O-PS alone (Δ*waaL*) had no significant effect on CPS retention **(Fig 1D)**. In contrast, loss of core-OS (Δ*wabG* and Δ*waaC*) significantly reduced the proportion of total CPS retained on the bacterial surface, by approximately two-fold relative to WT **(Fig 1D)**. This reduction was comparable between outer-core (Δ*wabG*; ∼1.8-fold) and inner-core (Δ*waaC*; ∼1.6-fold) mutants, indicating that core-LPS, but not O-PS, is required for WT-level CPS retention. Notably, loss of the core-PS also affected bacterial growth, as Δ*wabG* and Δ*waaC* showed significantly lower cumulative culture density after 16 hours of growth **(S1A Fig)**.

Similarly to the core-LPS mutants, Δ*wzi* showed significantly reduced (∼2-fold) CPS retention relative to WT **(Fig 1D)**. The Wzc_QK m_utant, however, exhibited an even more severe defect in CPS retention, as nearly the entire CPS was shed into the culture supernatant as cell-free EPS, which corresponds to a ∼6-fold reduction in CPS retention compared to WT **(Fig 1D)**. The reduced CPS retention in Δ*wzi* and Wzc_QK w_as further corroborated by higher levels of cell-free EPS observed upon staining of the isolated CPS and EPS fractions **(Fig 1E–F)**. Similar to the core-LPS mutants, Wzc_QK a_lso exhibited modest growth reduction, whereas the growth of Δ*wzi* was not affected **(S1A Fig)**. The observed reduction in CPS retention of Δ*wabG*, Δ*waaC* and Δ*wzi* were reversed by endogenous expression of the respective genes using a plasmid vector **(S2C Fig)**.

To test whether LPS core-OS and Wzi were similarly important for complete CPS retention in a classical *K. pneumoniae* strain, we determined CPS retention in *K. pneumoniae* strain MKP103. MKP103 is a carbapenem-susceptible derivative of the classical *K. pneumoniae* strain, KPN1H1, and expresses KL10-type CPS and O13-type LPS as determined by PathogenWatch (Kaptive v.3.1) [13,14,16]. Mirroring our observations in KPPR1, MKP103 transposon mutants of LPS core-OS (*wabG::cat* and *waaC::cat*) and wzi (*wzi::cat*) showed significantly reduced CPS retention compared to WT **(S3C Fig)**. Meanwhile, *waaL::cat* showed no change in CPS retention **(S3C Fig)**. Combined, these findings suggest that CPS biosynthesis proteins Wzi and Wzc, as well as the core-OS of LPS, are both broadly involved in retaining CPS at the surface of the outer membrane of *K. pneumoniae*.

### Divalent cations are important for complete CPS retention in *K. pneumoniae*

Metal cations are essential to bacterial cell physiology, ranging from acting as catalytic cofactors to maintaining membrane integrity [17,18]. For example, divalent cations stabilize LPS by forming salt bridges between anionic sugar moieties that would otherwise repel one another [18]. We hypothesized that divalent cations similarly stabilize LPS-mediated or Wzi-mediated CPS retention. To test this, we depleted Mg^2+^ and Ca^2+^ from the growth medium with ethylenediaminetetraacetic acid (EDTA) and measured CPS retention in WT, Δ*waaC* and Δ*wzi*.

EDTA reduced total CPS abundance and CPS retention in WT in a concentration-dependent manner, indicating that divalent cations support both CPS synthesis and retention **(Fig 2A–B)**. Δ*waaC* showed a similar concentration-dependent reduction in CPS retention, suggesting the requirement of divalent cations does not depend on intact core-LPS **(Fig 2B)**. In contrast, CPS synthesis and retention in Δ*wzi* mutant were not perturbed by EDTA treatment **(Fig 2A–B)**.

**Fig 2:**
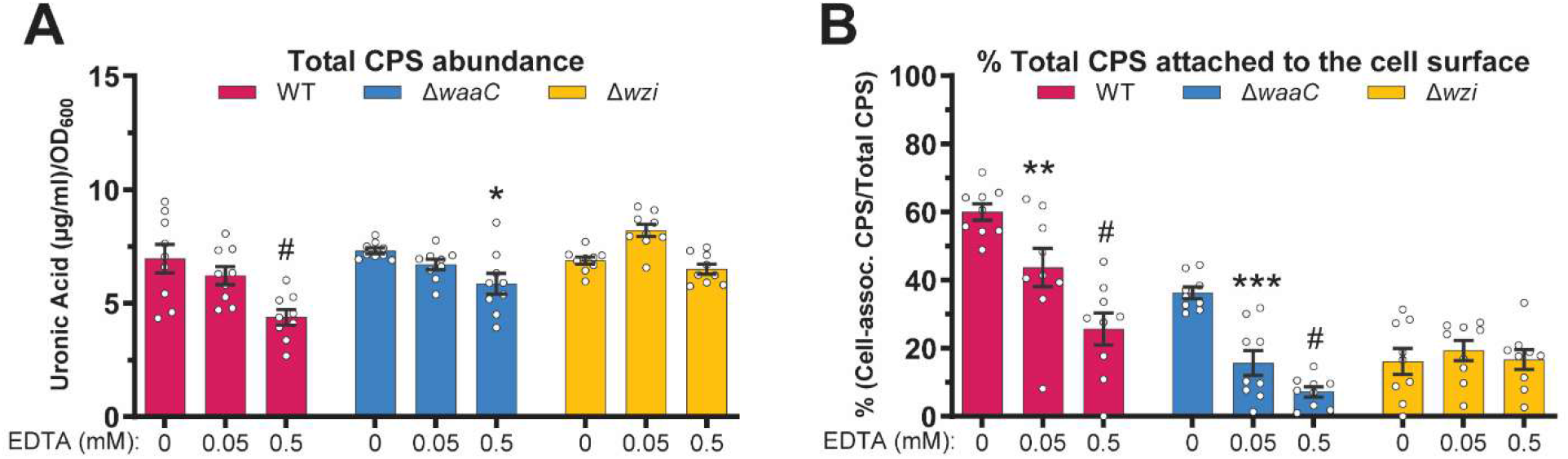
Divalent cations maintain stable retention of CPS on *K. pneumoniae* surface. *K. pneumoniae* strain KPPR1 WT, Δ*waaC* and Δ*wzi* were cultured in LB without or with 0.05 mM and 0.5 mM EDTA. **(A)** Total CPS abundance was determined by measuring uronic acid content in total crude CPS extracts and normalized to OD_600._ **(B)** The percentage of total CPS retained on the cell surface was calculated as the ratio of cell-associated CPS to total CPS measured in (A). Exact cell-associated CPS and cell-free EPS uronic acid levels are presented in S1A Fig. Data presented are the mean, and error bars represent the standard error of the mean. Statistical significance was determined for WT and each mutant by comparing the respective EDTA-added conditions (0.05 mM or 0.5 mM) to EDTA-free condition (0 mM) using two-way ANOVA with Šídák multiple-comparisons test. *P*-values are displayed above each bar; * indicates *p* ≤ 0.05, ** indicates *p* ≤ 0.01,*** indicates *p* ≤ 0.001, # indicates *p* ≤ 0.0001. Experiments were performed three independent times, each in biological triplicate (*n* = 9).

Next, we performed chemical complementation by supplementing EDTA-treated medium with MgSO_4 o_r CaCl_2._ Supplementing the EDTA-treated medium with either cation fully restored CPS retention in WT **(S4A–B Fig)**. In Δ*waaC*, CaCl_2,_ but not MgSO_4,_ partially restored CPS retention but this restoration did not reach statistical significance **(S4C–D Fig)**. In Δ*wzi,* CPS retention remained unchanged regardless of EDTA treatment or cation supplementation **(S4E–F Fig)**. Together, these results indicate that divalent cations are important for normal CPS synthesis and retention and that this requirement may be more specific to Wzi-mediated CPS retention. However, because a Δ*wzi* mutant exhibits a more severe CPS retention defect than Δ*waaC*, any additional contribution of cations to potential LPS-CPS interactions in Δ*wzi* may have been too small to be detected in this background. We therefore cannot completely rule out a role for divalent cations in stabilizing LPS-CPS interactions.

### Wzi extracellular loops may provide surfaces for capsular polysaccharide anchoring

Wzi is an outer-membrane protein encoded within the CPS biosynthesis loci of Gram-negative bacteria that employ Wzy-dependent CPS synthesis systems [9]. Its homologs have been identified in several enteric bacterial species including *Escherichia coli*, *Acinetobacter baumannii* and *Pseudomonas aeruginosa* [19–21]. Given the high sequence and structural similarity between *K. pneumoniae* and *E. coli* Wzi, much of our understanding of *K. pneumoniae* Wzi has been inferred from studies on *E. coli* Wzi. Studies on *K. pneumoniae*, *A. baumannii* and *E. coli* have established the role of Wzi in CPS anchoring [9,10,20]. However, despite the established role, contributions of individual structural elements of Wzi to CPS retention and the molecular basis of Wzi-mediated CPS retention remain poorly understood.

Structurally, Wzi comprises nine extracellular loops (L1–L9), eight periplasmic turns (T1–T8), a bundle of three periplasmic alpha helices (H1–H3) and 18 anti-parallel β-sheets that form the transmembrane region **(Fig 3A–B)**. Based on AlphaFold3-predicted structure, the extracellular face of Wzi is formed almost entirely by the extracellular loops, except L3, which is inwardly orientated **(Fig 3B)**. The extracellular surface formed by these loops display distinct electronegative and electropositive patches **(Fig 3C)**. Specifically, regions composed of L1 and L9 appear highly electronegative, while L2 and L7 are more electropositive than the surrounding loop surfaces **(Fig 3C)**. Anionic *K. pneumoniae* CPS may be retained by Wzi either through direct electrostatic attraction to electropositive surface patches or through cation-mediated bridging between anionic CPS and electronegative surface patches.

**Fig 3:**
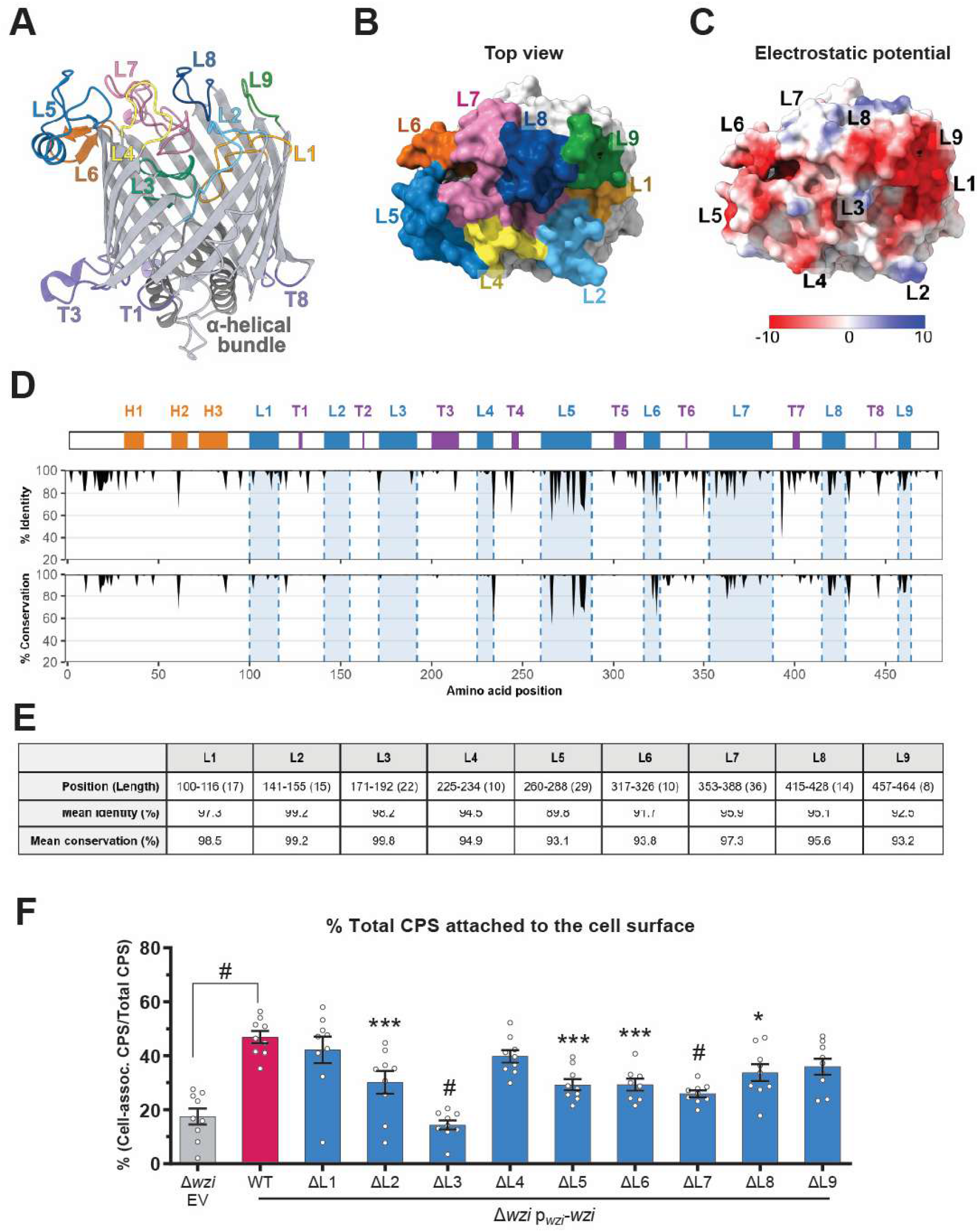
KPPR1 Wzi structure and conservation of sequences across *K. pneumoniae* strains. **(A)** Structure of KPPR1 Wzi (NCBI Protein Accession: WP_004180525) as predicted by AlphaFold-3. The extracellular loops (L1–L9), periplasmic loops (T1–T8) and the α-helical bundle (H1–H3; dark gray) are highlighted. **(B)** Top view (extracellular face) of the space-filling model of Wzi. The surfaces corresponding to each extracellular loop are highlighted in different colors. **(C)** The top view of Wzi showing distribution of electrostatic potential. Red indicates electronegative residues while blue indicates electropositive residues. **(D)** Conservation of Wzi residues across 122 reference *wzi* loci, including 110 unique KL-types. ‘% identity’ indicates frequency of the same amino acid at a given position, whereas ‘% conservation’ presents frequency of amino acids with similar physicochemical properties. The y-axis was confined to 20–100% to improve clarity. **(E)** The table presents the mean ‘% identity’ and ‘% conservation’ for each Wzi loop. **(F)** KPPR1 Δ*wzi* with empty vector (EV) or vector expressing WT or loop-substituted *wzi* (p*_wzi_*_-_*wzi*) were incubated in LB and total CPS abundance and cell-free abundance were measured by uronic acid quantification. The percentage of total CPS retained on the cell surface was calculated as the ratio of cell-associated CPS to total CPS measured. The corresponding total CPS, cell-associated CPS and cell-free EPS abundance are presented in S3C Fig. Data presented are the mean, and error bars represent the standard error of the mean. Statistical significance was determined by comparing strains expressing loop-substituted *wzi* or EV to the strain expressing WT *wzi*, using one-way ANOVA with Dunnett’s post-hoc test. *P*-values are displayed above each bar; * indicates *p* ≤ 0.05, *** indicates *p* ≤ 0.001, # indicates *p* ≤ 0.0001. Experiments were performed three independent times, each with biological triplicate (*n = 9*).

To investigate whether sequence variation across Wzi loops might contribute to CPS retention specificity, we analyzed 122 Wzi amino acid sequences representing 110 unique KL-types, with primary focus on the extracellular loops. These curated reference sequences have been previously used to develop a sequence-based typing scheme of *Klebsiella* capsule synthesis loci [22]. Overall, the Wzi sequence is largely conserved across strains with most regions showing over 80% identical residues (% identity) and above 90% of biochemically similar residues (% conservation). Despite this high degree of conservation, variations are evident and mostly concentrated on the region corresponding to the extracellular loops, particularly L5–L9 **(Fig 3D)**. As KL-types directly correlate with unique serotypes, sequence variation observed in the extracellular loop regions likely reflects a requirement for Wzi to accommodate structurally diverse CPS serotypes. Altogether, the structural organization and sequence diversity of the extracellular loops suggest that the extracellular surface is uniquely positioned to recognized and interact with CPS chains emerging from the Wza-Wzc export complex.

To systematically interrogate the function of extracellular loops, we constructed a series of Wzi complementation vectors (p*_wzi_*_-_*wzi*) in which each loop was individually substituted with a short, flexible glycine linker. Each loop-substituted variant was expressed in the Δ*wzi* background and compared for its CPS retention against vectors expressing WT *wzi* (Δ*wzi*-WT) or an empty vector control (Δ*wzi*-EV). All loop-substituted mutants, except for ΔL3, showed similar levels of Wzi protein expression compared to Δ*wzi*-WT **(S5A Fig)**. Expressing WT Wzi restored CPS retention in Δ*wzi* compared to Δ*wzi*-EV; although total CPS abundance in Δ*wzi*-WT remained lower than in Δ*wzi*-EV **(Fig 3F** and **S5B Fig)**. This result confirms that the loss of CPS anchoring in Δ*wzi* is attributable to Wzi function **(Fig 1D)**. Furthermore, all loop-substituted variants (Δ*wzi*-ΔL1–9) showed similar levels of total CPS abundance as Δ*wzi*-WT, except ΔL2 and ΔL3, which showed slightly higher total CPS abundance **(S5B Fig)**. Despite comparable total CPS abundance, substitution of L2, L3 and L5–L7 resulted in full or partial loss of CPS retention, whereas L1, L4 and L8–9 were functionally dispensable **(Fig 3F)**. Among the required loops, substitution of L3 had the most drastic effect, exhibiting CPS retention levels identical to Δ*wzi*-EV **(Fig 3F)**. Notably, L3 is folded inward and buried beneath the extracellular surface of Wzi based on the predicted structure **(Fig 3B)**. This suggests that the functional importance of L3 may lie in either proper folding of Wzi overall or providing optimal flexibility to the cleft formed by other loops, rather than directly interacting with CPS polymers **(Fig 3B)** [9]. The low level of Wzi-ΔL3 expression further supports the proposition that L3 may be important for proper protein folding **(S5A Fig)**. Among the functionally required loops, some regions had overall high sequence conservation (L2 and L3; >99%) and some regions carried greater positively-charged residue density (L2, L5 and L7) relative to the other loops **(Fig 3D–E)**. These findings collectively suggest that extracellular Wzi loops form a CPS retention site. We propose that, while some conserved loops (*e.g.*, L2 and L3) may be broadly required for maintaining Wzi structural flexibility, other less-conserved, positively-charged loops may contribute to electrostatic interactions with specificity to distinct K-types. However, the precise nature of the Wzi loop-CPS interaction remains speculative and will require further structural validation.

### CPS anchoring and LPS act at distinct stages of the complement cascade to confer serum resistance

Numerous reports in *K. pneumoniae* have demonstrated the protective roles of CPS and LPS against serum-mediated killing, otherwise known as serum resistance [23–25]. Bloodstream survival is important for dissemination from primary sites or hematological spread following bacteremia [26–28]. Whether disrupted CPS retention specifically protects against serum killing remains largely unknown. Therefore, we tested the survival of our CPS attachment mutants, including strains with truncated LPS and impaired CPS retention (Δ*waaC*) and strains with intact LPS but impaired CPS retention (Δ*wzi* and Wzc_QK)_, following incubation in 50% pooled normal human serum for 1 hour at 37°C. All mutants showed significantly reduced survival in human serum compared to WT, which maintained 24% survival of the total input **(Fig 4A)**. Specifically, all mutants, except Δ*waaC,* had less than 1% survival rate **(Fig 4A)**. The core-LPS mutant with partial CPS retention (Δ*waaC*), showed a mean survival rate of ∼10%, which was still significantly lower than WT **(Fig 4A)**. The increased susceptibility to human serum across all mutants was dependent on complement proteins, as survival of the mutants in the heat-inactivated serum was similar to WT **(Fig 4B)**.

**Fig 4:**
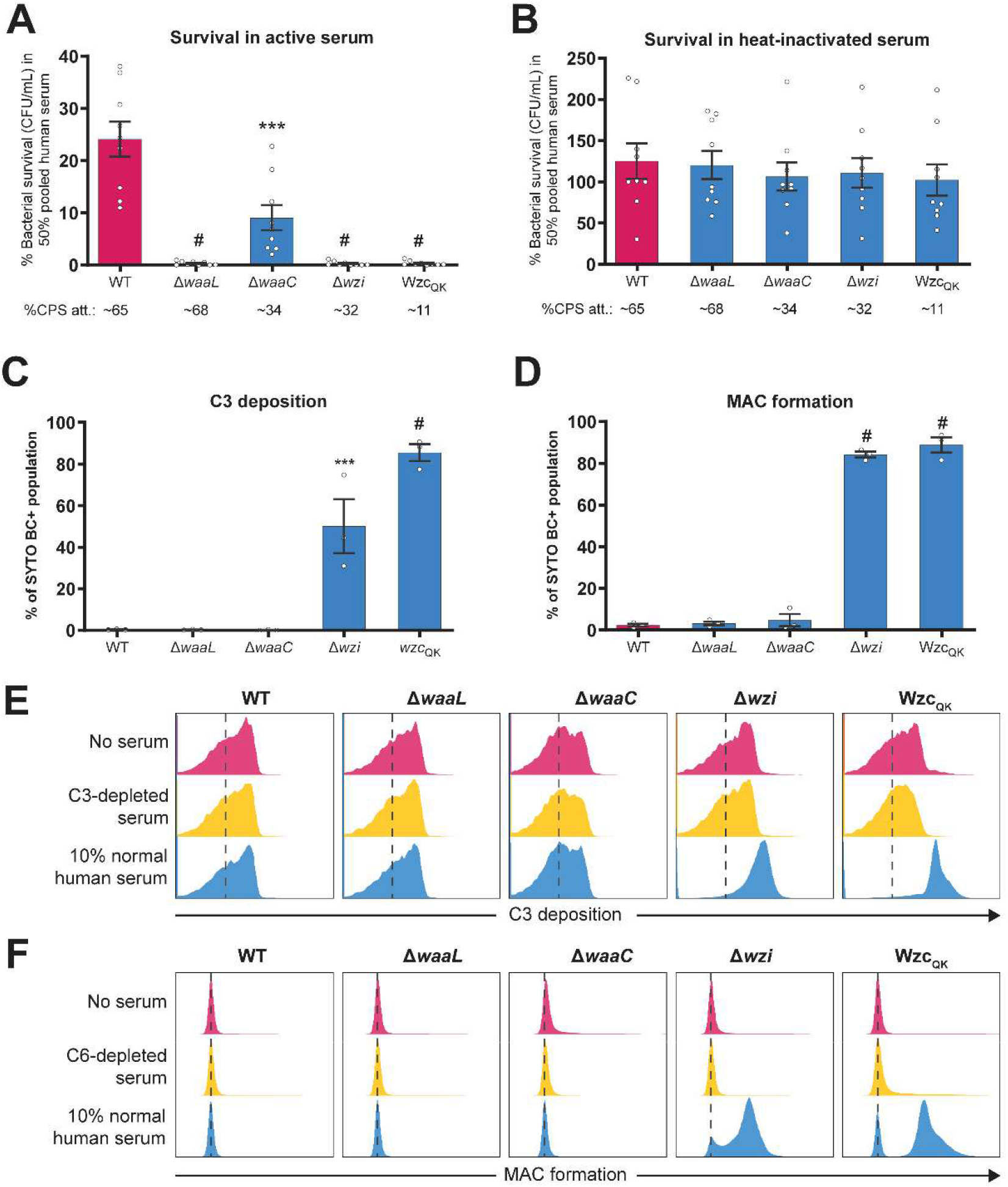
CPS and LPS both contribute to serum resistance, but increased complement deposition is associated more specifically with impaired CPS retention. KPPR1 WT and mutants were incubated with normal human serum and measured for resistance against serum-mediated killing and complement deposition. Bacterial survival was measured after incubating the WT and mutants at 37°C for 1 hour with **(A)** 50% normal human serum or **(B)** 50% heat-treated serum. Following incubation, bacterial survival was determined by enumerating CFUs and presented as the percent bacterial output relative to input. In a similar setup, bacteria were incubated with 10% normal human serum for 30 minutes and measured for **(C)** C3b deposition and **(D)** C5b-9 (membrane attack complex; MAC) formation using flow-cytometry. Syto BC+ is a nucleic acid stain used to select the total bacterial population and the y-axis in (C–D) represents percentage of the total Syto BC+ population. Representative flow-cytometry histogram of **(E)** C3 deposition and **(F)** C5b-9 (MAC) formation for data presented in (B–C). Bars represent the mean of 3 independent experiments performed in (A–B) triplicate (*n=9*) or (C–D) single replicate (*n=3*), and error bars represent the standard error of the mean. Statistical significance was determined using two-way ANOVA with Dunnett’s post-hoc test. Statistical significance was calculated by comparing mutants to WT. *P*-values are displayed above each comparison, *** *p* ≤ 0.001, # *p* ≤ 0.0001.

The complement system is a major innate immune defense mechanism and a critical component of serum-mediated defense against bacteria [29,30]. It promotes bacterial clearance through phagocytosis by labelling bacterial surfaces with opsonins and mediates direct bacterial killing by the formation of membrane attack complexes (MAC). The complement cascade involves more than 20 proteins and can be activated by three pathways that converge on the production of C3b, ultimately leading to the formation of MAC (C5b-9) on target surfaces, resulting in bacterial lysis [29,31]. Therefore, we hypothesized that the reduced serum survival amongst the mutants was due to increased complement deposition, particularly C3b and subsequent MAC (C5b-9) formation. To test this, we measured C3b deposition and MAC formation by flow-cytometry using C3b- and C5b-9-specific antibodies, after 30 minutes of bacterial incubation in 10% normal human serum [32]. Surprisingly, complement deposition across the mutants was not always consistent with their respective serum survival **(Fig 4C–F)**. Only the CPS-attachment mutants with intact LPS (Δ*wzi* and Wzc_QK)_ had increased C3b deposition and MAC formation relative to WT at the 30-minutes timepoint **(Fig 4C–F)**. Specifically, 50% and 85% of the Δ*wzi* and Wzc_QK b_acterial populations respectively, had C3b deposited on their surface, while both had MAC formation on over 84% of the population **(Fig 4C–D)**. This was significantly higher than serum-resistant WT, in which, less than 2% of the population showed both C3b deposition and MAC formation on its surface **(Fig 4C–D)**. In contrast, despite similarly lower serum susceptibility as Δ*wzi* and Wzc_QK,_ LPS truncation mutants Δ*waaL* (with normal CPS retention) and Δ*waaC* (with impaired CPS retention) had <1% C3b deposition and <4% MAC formation **(Fig 4C–D)**.

The discordance between serum survival and complement deposition indicates that the increased serum susceptibility of the LPS-truncation and CPS-attachment mutants is not always explained by increased complement deposition. Although fully (Δ*waaL*) or partially (Δ*waaC*) retained CPS chains may physically shield C3b deposition on the membrane, absence of O-PS in these mutants may affect steps downstream of C3b deposition sufficient to cause effective bacterial lysis despite minimal complement deposition. LPS truncation may allow for rapid MAC assembly and lysis, such that bacterial population that underwent lysis by the 30-minute timepoint are not represented in the analyzed population, causing an underestimation of true MAC formation. Alternatively, loss of O-PS may enhance the efficiency of C9 polymerization and membrane insertion even from the minimal complement deposited on the membrane, thus exhibiting enhanced serum-susceptibility with low C3b deposition and MAC deposition. This is consistent with a prior report that O-PS hinders MAC polymerization and insertion [25]. Overall, these data support that both LPS and CPS surface polysaccharides are critical for serum resistance in *K. pneumoniae*. Specifically, CPS protects by primarily limiting complement deposition directly on the membrane and LPS protects against complement-mediated lysis independent of complement deposition levels.

### CPS retention promotes bacterial hepatic and splenic fitness in a murine bacteremia model

To assess whether loss of CPS retention has any significant impact *in vivo*, competitive infection assays were performed using a murine bacteremia model. In this model, bacteria are directly inoculated into the bloodstream via tail vein injection (TVI), bypassing initial infection and dissemination, which usually precede Gram-negative bloodstream infections [26]. C57BL/6 mice were infected with an equal amount of WT and mutant in a combined total bacterial load of 2x10^5^ CFU/mL. Following infection, bacterial burdens in the blood and primary blood filtering organs, the liver, spleen and kidneys, were determined. All mutants were significantly outcompeted by WT in both liver and spleen **(Fig 5A–B)**. The overall competitive index (CI) was consistent across both organs, except for Wzc_QK,_ which showed a greater fitness defect in the spleen than in the liver (mean log_10 C_I: -1.75 *vs* - 1.06) **(Fig 5A–B)**. While the difference in CI between Δ*waaL* (partial LPS loss, fully retained CPS attachment) and Δ*waaC* (truncated LPS, decreased CPS attachment) was not statistically significant, our data suggest that a Δ*waaC* mutant exhibits a greater fitness defect than the Δ*waaL* mutant in both liver (Δ*waaC* vs Δ*waaL* mean log_10 C_I: -1.28 *vs* - 0.79) and spleen (Δ*waaC* vs Δ*waaL* mean log_10 C_I: -1.54 *vs* -1.08) **(Fig 5A–B)**. Blood and kidney samples were also collected from the infected animals, but they did not yield enumerable CFU for analysis. Altogether, these results indicate that Wzi- and LPS-mediated CPS retention are important for bacterial fitness during bloodstream infection, particularly for colonization of the liver and spleen.

**Fig 5:**
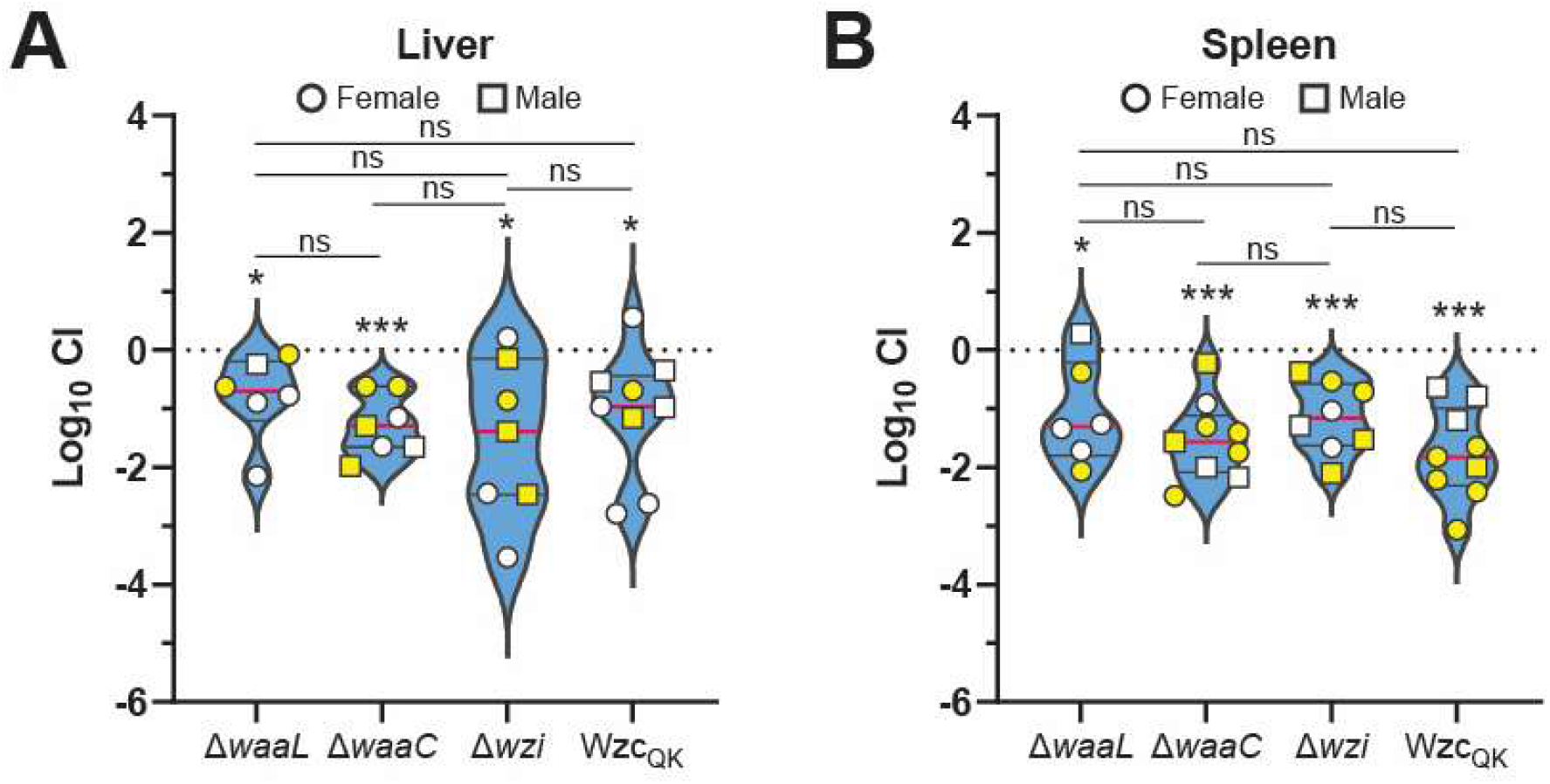
CPS and LPS are important determinants of hepatic and spleenic fitness of *K. pneumoniae*. Relative fitness of KPPR1 WT and the isogenic CPS attachment mutants were measured using a murine bloodstream infection model. A 1:1 mixture of KPPR1 WT and a mutant (combined total of 2x10^5^ CFU per mouse) were infected into C57BL/6 mice by tail vein injection. After 24 hours of infection, **(A)** Liver and **(B)** spleen were harvested. CFUs were enumerated using differential plating and log_10 c_ompetitive indices (CI; mutant:WT) were calculated using Equation 2. Each dot represents an individual mouse, where circular symbols are females, square symbols are males and either symbol filled with yellow color represents CFU recovery of either WT or mutant below the limit of detection. Data are presented as the median with interquartile range. Exact input and output CFU values are presented in S6 Fig. Statistical significance was determined using one-sample two-tailed *t* test and one-way ANOVA with Holm-Šídák multiple-comparisons correction. *P-*values displayed above each bar represent comparison of log_10 c_ompetitive index of a mutant to a hypothetical value of ‘0’. * indicates *p* ≤ 0.05, *** indicates *p* ≤ 0.001. Experiments were performed two independent times and included a total *n* per group of 6 (Δ*waaL*), 9 (Δ*waaC*), 8 (Δ*wzi*) and 9 (Wzc_QK)_.

## 3. DISCUSSION

Common bacterial CPS biosynthesis mechanisms like Wzy-dependent system have been extensively characterized. Yet, how these synthesized polysaccharides are retained on the bacterial cell surface and how differences in CPS retention due to inherent genetic changes or environment encountered in a host could affect bacterial fitness and pathogenesis remain poorly understood. Using a hypervirulent *K. pneumoniae* strain expressing the clinically relevant K2 capsule, we sought to identify and characterize the bacterial factors required for CPS retention, and determine whether disruption in proper CPS retention influences bloodstream infection [33]. Collectively, our findings indicate that CPS retention in *K. pneumoniae*, at least in the K2-capsule-type strain tested here, is a multifactorial process involving bacterial cell surface components including Wzi and LPS. Importantly, our findings further suggest that the ability to appropriately retain CPS may be as important as the ability to synthesize CPS itself for optimal bloodstream fitness.

Based on the literature, we selected the LPS O-PS, the LPS core-OS, Wzi and Wzc to interrogate their importance in CPS retention in our model *K. pneumoniae* strain [10,34]. Previous works in *K. pneumoniae* and *E. coli* have implicated LPS O-PS, LPS core-OS or both in anchoring CPS [10,35]. Contrary to what has been reported for the *K. pneumoniae* B5505 and NTUH-K2044 strains, the LPS O-PS was dispensable for CPS retention in our KPPR1 and MKP103 test strains **(Fig 1D** and **S3C Fig)** [10,36]. Genomic analysis using Kaptive revealed that B5505, NTUH-K2044, KPPR1 and MKP103 express distinct O-PS, O3γ, O1αβ,2α, O1αβ,2β and O13 respectively and synthesize distinct capsule types, K21 (B5505), K1 (NTUH-K2044), K2 (KPPR1) and K107 (MKP103). Thus, it is plausible that contribution of O-PS to CPS retention may be dependent on the specific combination of O- and K-types or the background of the strain. Among the other bacterial surface factors tested, LPS core-OS and Wzi showed significant role in CPS retention. Truncation of LPS to remove the core-OS (Δ*wabG* and Δ*waaC*) partially reduced CPS retention to about 40% of the total CPS, and this reduction was even more pronounced in Δ*wzi* (32%), compared to the 66% CPS retention in WT **(Fig 1D)**. On contrary, *wabG* and *waaC* disruption resulted in a greater reduction in CPS retention (20% retention) than *wzi* transposon (40% retention) **(S3C Fig)**. This finding agrees with several previous reports that truncation of core-OS from LPS significantly reduces the amount of CPS retained on the cell surface, while further suggesting that core-OS-mediated CPS retention may have a broadly conserved retention mechanism in *K. pneumoniae* [34,37–40].

Relative to the LPS O-PS, the LPS core-OS is highly conserved across Enterobacteriaceae, with *K. pneumoniae* expressing either type 1 or 2 core-OS [37,41]. Core-OS can be divided into outer-(distal to lipid-A) and inner-core (proximal to lipid-A) regions **(Fig 1A)**. Here, truncation of just the outer-core (Δ*wabG*) resulted in a similar reduction in CPS as the truncation of entire core-OS (Δ*waaC*) **(Fig 1D)**. This suggests that structural features contained in the outer core alone are sufficient to contribute to CPS retention in our test strain. Depending on the *K. pneumoniae* strain, core-OS consists of one to three galacturonic acid (GalA) residues, which contributes to the overall negative charge of the LPS polysaccharide component, particularly in the absence of phosphoryl groups found in core-OS of other Enterobacteriaceae members [39,41]. One of these GalA residues is found in the outer-core of both type 1 and 2 core-OS. These GalA units stabilize cell surface LPS organization via divalent cation-mediated interactions between adjacent LPS units [42,43]. A similar ionic interaction between LPS and CPS could therefore contribute to CPS retention in *K. pneumoniae*. A previous study support this hypothesis, where LPS and CPS from three different K2 strains co-eluted and then dissociated by chromatography with the metal chelator, EDTA [34]. Consistent with this model, we observed decreased CPS retention with increasing EDTA concentrations and restored CPS retention upon Ca^2+^ and Mg^2+^ supplementation in the growth medium, further supporting a role for divalent cations in CPS retention **(Fig 2B** and **S4 Fig)**. However, since Ca^2+^ and Mg^2+^ are broadly involved in outer-membrane stability, our experimental setup does not conclusively distinguish between the direct and indirect role of divalent cations in LPS-CPS or Wzi-CPS interactions from its indirect effect on cell-membrane integrity and ensuing impact on CPS retention [43]. Nevertheless, the data presented here, along with prior work, support that divalent metal ion bridges may facilitate LPS-CPS and Wzi-CPS electrostatic interactions, contributing to CPS retention on the bacterial outer-membrane.

In line with observations from *E. coli* and other *K. pneumoniae* strains, Wzi emerged as an important CPS retention factor in KPPR1 [9,10]. To define the mechanism by which Wzi supports CPS attachment, we interrogated the involvement of the extracellular-facing loop domains that are positionally poised to facilitate interactions with CPS chains following export. The extracellular surface is primarily formed by nine extracellular loops **(Fig 3A–B)**. Electrostatic potential mapping shows that the surface is largely electronegative, except for a distinct electropositive patch including and surrounding L7 **(Fig 3C)**. In *E. coli*, the electropositive patch is thought to form electrostatic interaction with anionic CPS [9]. However, *E. coli* Wzi has more prominent electropositive patches than KPPR1 that nearly spans L4–L8 [9]. In *E. coli* K30, specifically, L3, L6 and L7 are required for CPS retention while L8 is dispensable [9]. To assess whether a similar electrostatic interaction could be broadly conserved across *K. pneumoniae* strains, we aligned Wzi sequences from over 120 distinct KL-type. We found revealed that Wzi is highly conserved across distinct KL-types, yet some sequence diversity was observed, which was concentrated primarily in the loop regions **(Fig 3D–E)**. Loops L3 and L7, that reportedly had functional importance in *E. coli*, were conserved across different *K. pneumoniae* Wzi with >97% sequence similarity [9]. Further, we demonstrated a role for these domains in CPS attachment through mutagenesis of individual Wzi loop domains, where each loop was replaced with five glycine residues that contains a net zero charge at physiological pH. In a series of KPPR1 Wzi loop substitution mutants, L2, L3 and L5, L6 and L7 were identified as important for Wzi-mediated CPS retention as these mutants exhibited significantly reduced CPS retention compared to Wzi-WT control **(Fig 3F)**. The observation with L3, L6 and L7 are consistent with the prior report from *E. coli* [9]. However, it should be noted that L2 and L5 were not tested for their function in the mentioned study. Of note, contrary to other loops, L3 is bent inwards towards the β-barrel and its substitution was accompanied with reduced Wzi expression, which is suggestive of L3’s role in stabilizing the Wzi conformation as well as providing flexibility to the CPS-binding “cleft” formed by other CPS-interacting loops. The functionally important loops identified in our study were not necessarily limited to just positively charged loops. Loops with primarily electronegative residues (L6), neutral residues (L3 and L5), and electropositive residues (e.g., L7 and L2) were found to be functionally important. Among these, L2, L3 and L7 are highly conserved (>97%) across distinct *K. pneumoniae* KL-types. Collectively, these findings suggest that the electropositive and highly conserved loop regions might provide a general site for electrostatic interaction to negatively charged CPS, while the variable and neutral to negatively charged loops may be responsible for accommodating more than >80 distinct CPS types found in *K. pneumoniae*.

Wzi has long been described as an outer-membrane lectin involved in CPS retention based on *in vitro* Wzi-CPS interaction assays [9]. Although *wzi* is a conserved locus encoded within the broader *K. pneumoniae* CPS biosynthesis locus, it is not considered an essential component of the CPS biosynthesis machinery. Intriguingly, however, we observed a moderate but significant reduction in total CPS abundance **(Fig 1C)** along with reduced molecular weight of ‘form C’ CPS, indicated by a downward shift of the corresponding polysaccharide band in SDS-PAGE **(Fig 1E)**. These findings raise the possibility that Wzi-dependent CPS organization and CPS biosynthesis are at least in part, functionally linked. Consistent with this possibility, we observed a near complete loss of CPS retention in KPPR1 expressing a Q395K variant of the CPS polymerization regulator Wzc. In comparison to WT, which retains about 66% of the total CPS on its surface, Wzc_QK s_train retains only ∼11% of the total CPS. Notably, one-third of the biological replicates showed complete loss of CPS retention **(Fig 1D)**. Wzc is an integral component of Wzy-dependent CPS biosynthesis systems and regulates CPS polymerization rate *i.e.*, CPS chain length [44]. It also interacts with the Wza OM pore complex, forming the CPS export machinery that transports newly synthesized CPS polymers from the periplasm to the cell surface [45]. Since Wzc is an inner-membrane protein, it is unlikely to anchor CPS directly. The Q395 residue is located in the periplasmic domain of Wzc which is important for its interaction with the outer-membrane export protein Wza. Thus, the Q395K substitution may have brought conformational changes that affect CPS polymerization activity while also impacting its transport through the Wzc-Wza export complex consequently affecting how the CPS chains are presented to Wzi. We did not detect any significant conformational change using AlphaFold in Wzc monomer with Q395K substitution though the possibility of conformational change cannot be ruled out. Nevertheless, our data indicate that the core biosynthetic steps determining CPS polymer properties, including chain length, polymerization rate and composition, may influence CPS surface retention. Together, the bidirectional impact between Wzi-dependent CPS retention and Wzc-dependent CPS synthesis suggest that CPS biosynthesis and CPS surface retention may have functional overlap to some extent. Further studies are needed to dissect the influence of CPS biosynthesis and retention on each other, some of which includes assessing whether Wzc_QK s_ynthesizes compositionally different CPS and the affinity of Wzi towards different forms of CPS (types A, B or C).

Overall, neither Wzi nor LPS core-PS accounted for complete CPS retention in KPPR1. Almost half of the total CPS that is retained by WT remained anchored to the surface in Δ*wzi*, while about 75% was still anchored in Δ*waaC*. This pointed to the possibility that complete CPS retention is a collectively mediated by LPS and Wzi. To test this, we attempted to generate a Δ*waaC*-Δ*wzi* double mutant without success. Truncation of LPS combined with the removal of the Wzi OM protein might lethally destabilize the membrane. Nevertheless, our findings with single mutants support a model where Wzi is a primary contributor to CPS retention, but other surface components, like LPS, are also required for proper CPS retention, thus making CPS retention in *K. pneumoniae* a truly ‘multifactorial process’.

Both CPS and LPS have been implicated in conferring resistance to complement-mediated killing in serum [24,25]. Our findings support this, as both LPS-truncation mutants and CPS-retention mutants exhibited significantly increased serum susceptibility **(Fig 4A–B)**. However, the contribution of LPS and CPS to complement resistance may depend on the O- and K-types of the tested strain, and the protective mechanisms observed here may not be generalizable across *K. pneumoniae* strains [25,46]. Notably, Δ*wzi* and Wzc_QK,_ which fail to properly retain CPS, showed increased C3b and MAC deposition that was consistent with their reduced serum survival **(Fig 4C–D)**. This finding supports our model that surface-attached CPS is important in protecting from serum-derived factors. In contrast, the serum survival of the LPS mutants was not consistent with complement deposition. Specifically, all three LPS mutants (Δ*waaL*, Δ*wabG* and Δ*waaC*) had low C3b and MAC deposition despite being similarly susceptible to serum as Δ*wzi* and Wzc_QK (_**Fig 4A–D)**. This discordance indicates that complement-resistance conferred by LPS, particularly O-PS, is distinct from CPS. Importantly, the amount of complement deposition does not necessarily predict the bactericidal effect of the terminal MAC [25]. A study of *K. pneumoniae* O1-type LPS demonstrated that loss of the O-PS resulted in reduced C3b and C5b-9 deposition despite increased complement-mediated killing [25]. In this instance, the O-PS mutant exhibited increased C9 polymerization and more efficient MAC insertion into the bacterial membrane [25]. This illustrates how relatively less complement deposition can still result in effective membrane damage. A similar mechanism could explain the observed discordance between serum sensitivity and MAC deposition observed in our LPS mutants. We therefore propose that CPS and LPS provide similar levels of protection against complement-mediated killing, although they may act at distinct stages of complement activity. The high-molecular weight CPS polymers (*i.e.*, longer chains) might be primarily responsible for preventing initial complement deposition on the surface, while the low molecular weight LPS (*i.e.*, shorter chains) may influence effective assembly of terminal MAC and membrane insertion. Further assessment of terminal MAC assembly including measurement of C9 polymerization and membrane insertion will be required to answer how LPS and CPS distinctly contribute to *K. pneumoniae* complement resistance.

Next, we assessed whether proper CPS retention impacts pathogenesis *in vivo* by determining bacterial fitness during bloodstream infection using a murine bacteremia model. Although attempts were made to enumerate bacteria from the blood and the primary blood filtering organs the liver, spleen and kidneys, we did not recover enumerable bacteria from the blood and kidneys. The lack of subsequent bacterial recovery from the blood is expected as *K. pneumoniae* is rapidly killed by blood factors and is a known limitation of the model used in the study [26]. All mutants impaired in CPS retention were consistently outcompeted by WT in both the liver and spleen, suggesting that proper CPS retention is important for optimal fitness in these systemic sites **(Fig 5A–B)**. The Δ*waaL* mutant, which exhibits proper CPS retention but lacks the O-PS, was also similarly outcompeted **(Fig 5A–B)**. While this result appears consistent with the important role of the O-PS in human serum survival described above, murine (C57B/L6) serum is ineffective at killing KPPR1, which is also reported for other Gram-negative species **(Fig 4A–B)** [47,48]. This suggests that the fitness defects observed in the LPS-truncation and CPS-retention mutants in our murine model may reflect impaired protection from cell-mediated immune clearance rather than complement-mediated killing. The TVI model used here bypasses the initial site colonization and introduces bacteria directly into the bloodstream [26]. Thus, this model primarily assesses the contribution of the bacterial surface polysaccharides after bacterial entry to the bloodstream, including survival and persistence in blood-filtering organs such as the liver and spleen. Upon entry into the vasculature, a surface-retained capsule may protect the bacteria from immediate immune-mediated clearance and facilitate subsequent survival within systemic sites. The fitness defect observed here in CPS-retention mutants is consistent with a prior study demonstrating that non-capsulated *K. pneumoniae* strains and strains expressing low-virulence capsule serotypes have reduced bloodstream fitness, more specifically due to increased capture and uptake by liver-resident macrophages (Kupffer cells) [49]. Thus, these findings suggest that CPS surface retention may enhance bacterial fitness at multiple stages of bloodstream infection. Interestingly, there was no clear distinction between LPS and CPS-retention mutants in terms of contribution of either polysaccharide to bloodstream fitness. Both groups of mutants were equally outcompeted by the WT and additionally, the Δ*waaC* mutant with truncated LPS and reduced CPS retention did not exhibit any greater defect than either Δ*waaL* (loss of O-PS) or Δ*wzi* (partial CPS retention loss) **(Fig 5A–B)**. Although a greater defect of Δ*waaC* was observed in the spleen, it did not reach statistical significance **(Fig 5B)**. This greater fitness defect of Δ*waaC* in the spleen compared to the liver may stem from niche-dependent differences in immune factors. Overall, our findings support the idea that CPS retention is an important determinant of *K. pneumoniae* fitness during bloodstream infection.

In summary, our findings suggest that CPS retention in *K. pneumoniae* is a multifactorial process that jointly involves both LPS and Wzi, and likely mediated by cation-mediated electrostatic interactions. Further, we report that CPS biosynthesis and retention process may be functionally linked to each other. Although the contribution of these CPS-retention components may vary with strain and CPS serotype, our findings indicate that the CPS-mediated protection depends on its proper retention and organization on the cell surface and not just its synthesis. Further elucidation of the molecular basis for CPS retention may open new avenues for identification of anti-virulence targets to tackle the emerging threat of hypervirulent and multidrug-resistant *K. pneumoniae*.

## 4. MATERIALS AND METHODS

All experimental procedures comply with the ethical guidelines established by the institution involved. Institutional Biosafety Committee (IBC) approvals were obtained from the University of Pittsburgh (PROTO202400096). This study was performed in strict accordance with the recommendations in the *Guide for the Care and Use of Laboratory Animals* [50]. The University of Pittsburgh Institutional Animal Care and Use Committee approved this study (protocol 24105753; PI: Laura Mike).

### Bacterial strains and culture conditions

All bacterial strains and plasmids used in the study are detailed in S3 Table. Bacterial strains were cultured in lysogeny broth (LB) (5 g/L yeast extract, 10 g/L tryptone, 0.5 g/L NaCl) and incubated aerobically at 37°C and 200 rpm shaking, unless otherwise noted. Solid culture medium was prepared by adding 20 g/L bacto-agar to LB [51]. When appropriate, antibiotics were added at the following concentrations: kanamycin (25 µg/mL), gentamicin (15 µg/mL) and chloramphenicol (20 µg/mL for *E. coli* or 80 µg/mL for *K. pneumoniae*).

### Uronic acid quantification

Capsular polysaccharide abundance was measured by quantifying uronic acid content [52,53]. In brief, capsular polysaccharide was purified by ethanol precipitation from 250 µL of overnight culture. To 250 µL of overnight bacterial culture, either 50 µL of 1% Zwittergent 3-14 or 50 µL of ultra-pure water was added to isolate total capsular polysaccharide or cell-free EPS, respectively. The mixture was precipitated at 17,000 *x g* for 5 mins and the polysaccharide was collected by precipitating 100 µL of the uppermost supernatant in 400 µL ice-cold ethanol. Then, uronic acid content in the purified polysaccharide was quantified by using a colorimetric assay involving 3-hydroxyphenyl phenol [52]. Cell-associated CPS uronic acid level was deduced by subtracting the uronic acid levels in cell-free EPS from the total capsular polysaccharide.

### Molecular cloning, transformation and sequencing

Oligonucleotides used for cloning in this study are listed in S4 Table. *K. pneumoniae*isogenic knockout mutants were constructed using the λ Red recombineering adapted to *K. pneumoniae* [54–56]. Complementation plasmids were constructed by PCR amplification of the plasmid backbone (pAYCY184Δ*tet* or pBBR1-MCS5) and gene fragments (native promoter and gene of interest). The gel-purified PCR products (Monarch, NEB) were then assembled using NEBuilder HiFi DNA Assembly Mix (NEB) for 1 hour at 50°C. The resulting ligation product was then transformed into TOP10 *E. coli* through heat-shock and confirmed via sequencing. The resulting plasmid was transformed into *K. pneumoniae* strains by electroporation [56]. Plasmid constructs were validated by sequencing (Plasmidsaurus).

### Bacterial growth assay

Bacterial growth curves were generated by measuring OD_600 a_t specific intervals [51]. Overnight cultures were setup in LB and incubated aerobically at 37°C and 200 rpm. Then, a 1:1000 subculture of the overnight culture was prepared in 3 mL LB and incubated till mid-log. One hundred µL of 0.001 OD_600 e_quivalent subculture was then transferred to a sterile 96-well plate. Additionally, 100 µL of sterile water was added to the outermost wells of the plate to prevent samples from drying out. Finally, OD_600 w_as measured every 30 min for 16 hours at 37°C and continuous orbital shaking (282 rpm) using a microplate reader (SYNERGY HTX, Agilent).

### CPS extraction and visualization

Cell-associated CPS and cell-free EPS were purified by ethanol precipitation and visualized using Alcian blue and silver stain [11,52]. Cell-attached CPS were purified from bacteria equivalent to 1.5 OD_600._ Cells were pelleted at 21,000 × *g* for 15 min to remove cell-free EPS. The bacterial pellet was resuspended in 1 mL of PBS and centrifuged at 21,000 × *g* for 15 min. All but 250 µL of the supernatant was removed, mixed with 50 µL 3-14 1% Zwittergent and incubated at 50°C for 20 min. The incubated mixture was then centrifuged at 21,000 × *g* for 5 min to collect the upper 100 µL of the supernatant. The supernatant was added to a tube containing 400 µL ice-cold ethanol to precipitate polysaccharide.

For staining, CPS preparations were mixed with 4 × SDS gel loading dye (3:1, sample:dye) and 20 µL of the mixtures were resolved using 4–15% TGX stain-free precast gel (Bio-Rad). The electrophoresis was performed for 4.5 hours at 300 V on ice at 4°C. After electrophoresis, the gel was stained with 0.1% alcian blue (0.1% wt/vol ThermoFisher Alcian Blue 8 Gx in stain base solution) for 1 h. It was then stained with Pierce Silver Stain Kit (ThermoFisher) following the manufacturer’s instructions.

### LPS extraction and visualization

LPS was extracted using iNtRON LPS extraction kit (iNtRON Biotechnology) from 5 mL of overnight culture adjusted to a final cell density of OD_600 =_ 1. Manufacturer’s instruction was followed with minimal modifications. Briefly, the cell density-adjusted cultures were centrifuged at 21,000 *x g* for 10 minutes at room temperature and pellet was resuspended in lysis buffer. It was then treated with chloroform followed by purification buffer to collect a clear supernatant. Then, the supernatant was mixed with 70% ethanol to precipitate out polysaccharide and centrifuged at 21,000 *x g* at 4°C to collect LPS pellet. The pellet was resuspended in 50 µL of 10 mM Tris-HCl (pH 8.0) by boiling at 100°C for 2 minutes and stored at -20°C until further use.

The LPS preparation was resuspended in 4X SDS gel loading dye (3:1, LPS:dye) and 20 µL of the mixtures were resolved using a 4–15% TGX stain-free precast gel (Bio-Rad). The electrophoresis was performed at 140 V with constant current at room temperature for ∼50 minutes. After electrophoresis, the gel was stained with Pierce Silver Stain Kit (ThermoFisher) following the manufacturer’s instructions.

### Wzi structure modeling and visualization, and alignment

Wzi amino acid sequence (WP_004100525.1) was used to model the structure of KPPR1 Wzi. AlphaFold 3 (https://www.alphafoldserver.com) was used for structure prediction, and UCSF ChimeraX V1.9 was used for visualization of the modelled structure [57].

For alignment of Wzi across unique KL-type, a previously published curated list of accession codes for each reference K-locus were used to retrieve the whole genome sequence [22]. Respective Wzi protein sequences were downloaded from NCBI and aligned to KPPR1 reference Wzi sequence using MAFFT.

### Western blotting

Western blotting of his-tagged Wzi was performed using the protocol described by Khadka and Ring et al. with the following modifications [11]. One milliliter of each overnight bacterial culture was pelleted at 21,000 *x g* for 15 minutes at 4°C and lysed with 250 µL lysis buffer by sonication (50% amplitude, 2 x 5s). The lysis buffer was prepared as follows: 7 mL BugBuster protein extraction reagent (MilliporeSigma; Cat no.: 70584-4) was mixed with one cOmplete Mini, EDTA-free Protease Inhibitor Cocktail tablet (Roche; Cat no.: 11836170001). Cell lysates were measured for total protein concentration by Pierce BCA Protein Assay kit (ThermoFisher; Cat no.: 23227) and adjusted to 1.4 µg/µL with 1x SDS-PAGE gel loading dye. Ten microliter of such prepared lysate was resolved by SDS-PAGE, transferred to a nitrocellulose membrane using TransBlot Turbo Transfer System (BioRad) and blocked with 10 mL of EveryBlot Blocking Buffer (BioRad; Cat no.: 12010020) for 1 hour. The blocked membrane was incubated with 1:5000 mouse anti-His-tag-HRP antibody (SouthernBiotech; Cat no.: 4603-05) for 1 hour, washed with 1x TBS-T buffer and developed using Pierce ECL Western Blotting Substrate (ThermoFisher; Cat no.: 32106). It was then imaged using the BioRad ChemiDoc XRS+ imaging system.

### Human serum survival assay

Bacterial survival in pooled human serum was measured as described by Mike et al. and Kinney et al [32,54]. One mL overnight bacterial culture was centrifuged at 16,249 × *g* for 10 min and resuspended in sterile PBS. It was then adjusted to OD_600 e_quivalent of 0.2 in 200 µL, followed by 1:100 dilution in sterile PBS to get a final bacterial inoculum of approximately 2×10^6^ CFU/mL. Then, 45 µL of the OD-adjusted bacterial preparations were incubated in 50% normal human serum (Innovative Research; Catalog no.: ISCER100ML) or heat-inactivated serum (inactivated at 58°C for 1 h) at 37°C for 60 min. Input and output bacterial load were determined by enumerating CFUs using serial dilution and plating on LB agar. Bacterial survival was presented as the percentage of input CFUs surviving after incubation.

### C3 and C5b-9 deposition assay

Complement deposition on bacterial surface was measured by flow cytometry as described by Bain et al. and Kinney et al [32,58]. Overnight bacterial cultures were sub-cultured (1:100) in LB and incubated at 37°C with 200 rpm shaking. After the bacterial culture entered exponential growth phase, the culture density was adjusted to OD_600 o_f 0.2. The cells were then washed and resuspended in GVB° (ComplementTech #B103; no Mg or Ca). Then, 250 µL of bacteria was incubated with 25 µL of MgEDTA along with either 5% C3-depleted serum (ComplementTech #A314), 5% C6-depleted serum (ComplementTech #A323) or 10% normal human serum (ComplementTech #NHS) for 30 mins at 37°C with periodic mixing. After incubation, the reaction was quenched using 750 µL cold sterile PBS, pelleted (1,900 × *g* for 5 min). It was followed by two successive washes of the cells using 500 µL PBS. Each sample was then resuspended in 100 µL 2% FBS in PBS. Then, 50 µL of the cells were treated with one or all the following antibodies (2.5 μL ; 1:20 dilution per 10^6^ cells): C3b/iC3b-APC (Biolegend #846106), C5b-9-Alexa Fluor 549 (Novus #05120AF594) and SYTO BC green nucleic acid stain (Invitrogen by Thermo Fisher Scientific #S34855). The reaction was incubated at 4°C for 30 min and then washed twice with PBS (centrifuged at 1,900 × *g* for 5 min, supernatant removed, then resuspended in 500 μL PBS). Samples were analyzed on a Fortessa 7 cytometer (thresholds: 200 for FSC and 200 for SSC).

### Murine model of infection

Eight to 12-week old C57BL/6 mice (Charles River, strain 000664 RB07) were infected in the bacteremia model. Briefly, KPPR1 WT and mutant strains were cultured overnight from single colonies in LB at 37°C. The following day, mice were inoculated at a 1:1 (WT:mutant) ratio by TVI. The inoculum density was adjusted to a final concentration of 2×10^6^ CFU/mL in sterile saline. The exact input ratio was confirmed by differential plating.

Mice were inoculated by TVI with 100 µL of the inoculum (*i.e.*, 2×10^5^ CFU) per mouse. At 24 hours post-inoculation, the mice were humanely euthanized and blood (by cardiac puncture), liver, kidneys and spleen were collected and homogenized in sterile PBS and plated on LB agar or LB agar containing the appropriate antibiotics. CFUs were enumerated after overnight incubation at room temperature. The competitive index (CI) was calculated using the formula shown in Equation (2).

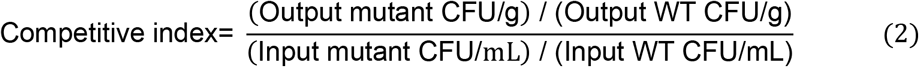

The limit of detection for *K. pneumoniae* detection in these experiments was 100 CFU/mL. For all samples below the limit of detection, we assumed a bacterial concentration of 99 CFU/mL. Organ samples with total bacterial count (WT+mutant) plated on LB showing CFU below LOD were excluded from analysis. Additionally, organ samples where total bacterial CFU (WT+mutant) was lower than that of mutant CFU were also excluded.

### Statistics

All replicates represent biological replicates and were performed at least three times independently, unless otherwise noted. Statistical analyses were computed in Graphpad Prism 10 (v10.5.0; GraphPad Software, LLC). Flow cytometry data were analyzed and plotted using FlowJo 11 (v11.2.0; Becton, Dickinson and Company). For experiments comparing more than two groups on two independent variables, significance was calculated using two-way ANOVA with Dunnett’s post-hoc test or Šídák correction. Meanwhile, one-way ANOVA with either Dunnett’s post-hoc test or Šídák correction was applied to calculate significance between more than two groups on one independent variable. Statistical differences between two groups with independent variables were calculated using student’s t-test. In all instances, results were considered statistically significant if the *P*-value was less than or equal to 0.05. Unless otherwise noted, the graphs represent an average of replicates with standard error of mean.

## ACKNOWLEDGEMENTS

Research reported in this publication was supported by the University of Toledo, University of Pittsburgh, 23CDA1056712 (L.A.M.), 24PRE1197026 (S.K.), 26PRE1561982 (E.L.K) from the American Heart Association, and K22 AI145849 (L.A.M.), R35 GM150588 (L.A.M.) and TL1 DK143271 (Z.J.R.) from the National Institutes of Health. This content is solely the responsibility of the authors and does not necessarily represent the official views of the funding agencies.

We thank members of the Mike Lab, Program in Microbiology and Immunology and Division of Infectious Diseases at the University of Pittsburgh for sharing technical resources, and valuable guidance and feedback.

---

**S1 Fig:**
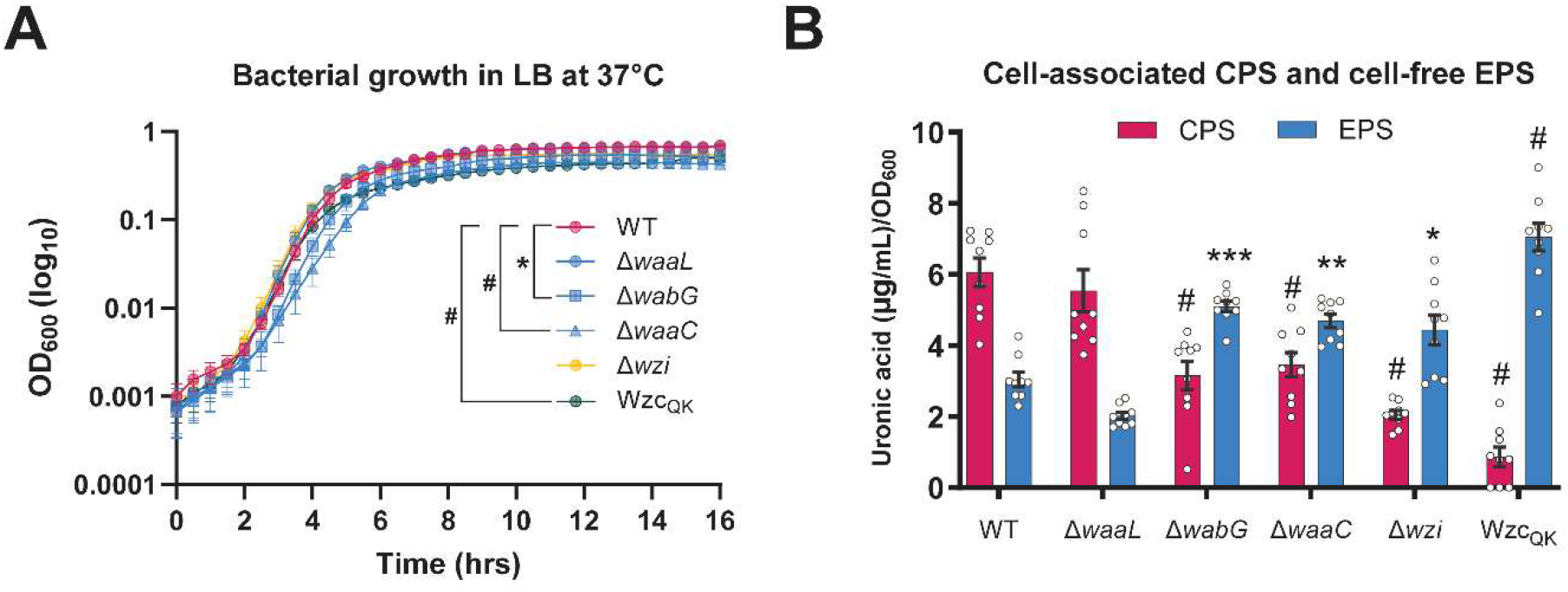
Deletion of CPS retention components increases proportion of cell-free EPS without impairing *K. pneumoniae* growth *in vitro*. KPPR1 WT and the mutants were cultured in LB and their cell-associated CPS, cell-free EPS abundance and growth kinetics were analyzed. **(A)** Cell-associated CPS and cell-free EPS abundance were determined by uronic acid quantification. Cell-associated CPS was calculated by subtracting cell-free EPS from total CPS abundance. The corresponding ‘total CPS abundance’ and ‘% total CPS attached to the cell surface’ are presented in Fig 1C and 19D, respectively. **(B)** Bacterial growth curves were generated by measuring OD_600 o_f WT and the mutants every 30 minutes cultured aerobically for 16 hours at 37°C. (A,B) Data presented are the mean, and error bars represent the standard error of the mean. Statistical significance was determined by comparing each mutant to WT using (A) two-way ANOVA with Dunnett’s post-hoc test or (B) one-way ANOVA with Dunnett’s post-hoc test. *P*-values are displayed (A) above the bars or (B) alongside data legend with vertical brackets, for each comparison; * indicates *p* ≤ 0.05, *** indicates *p* ≤ 0.001, # *p* indicates ≤ 0.0001. Experiments were performed three independent times, each in biological triplicate (*n* = 9).

**S2 Fig:**
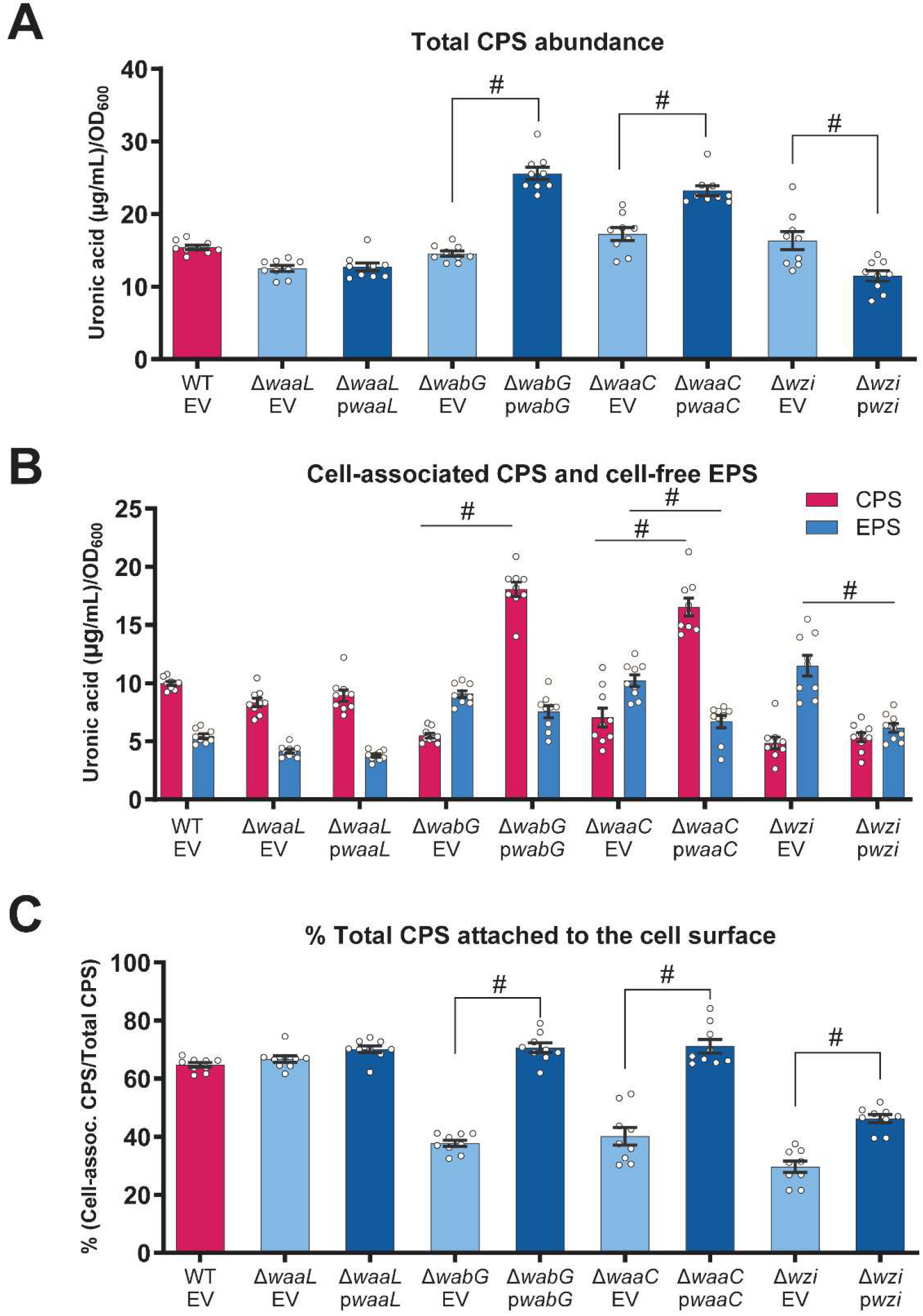
Genetic complementation of the LPS and *wzi* deletion strains. KPPR1 mutants Δ*waaL*, Δ*wabG*, Δ*waaC* and Δ*wzi* were genetically complemented using the pBBR1-MCS5 vector carrying the respective endogenous promoter and gene. The strains were cultured in LB and their cell-associated CPS and cell-free EPS abundance were analyzed. **(A)** Total CPS abundance was determined by measuring uronic acid content in total crude CPS extracts and normalized to OD_600._ **(B)** Cell-associated CPS and cell-free EPS abundance were determined by uronic acid quantification. Cell-associated CPS was calculated by subtracting cell-free EPS from total CPS abundance. **(C)** The percentage of total CPS retained on the bacterial cell surface was calculated as the ratio of cell-associated CPS to total CPS shown in (B). Data presented are the mean, and error bars represent the standard error of the mean. Statistical significance was determined by comparing adjacent empty vector (EV) and complementation strains to WT using two-way ANOVA with Šídák correction for multiple comparisons. *P*-values are displayed above the bars with connecting lines; * indicates *p* ≤ 0.05, ** indicates *p* ≤ 0.01, # indicates *p* ≤ 0.0001. Experiments were performed three independent times, each in biological triplicate (*n* = 9). ‘EV’ indicates strain with an empty pBBR1 vector and p*gene* indicates strain with pBBR1 encoding the gene under its respective endogenous promoter.

**S3 Fig:**
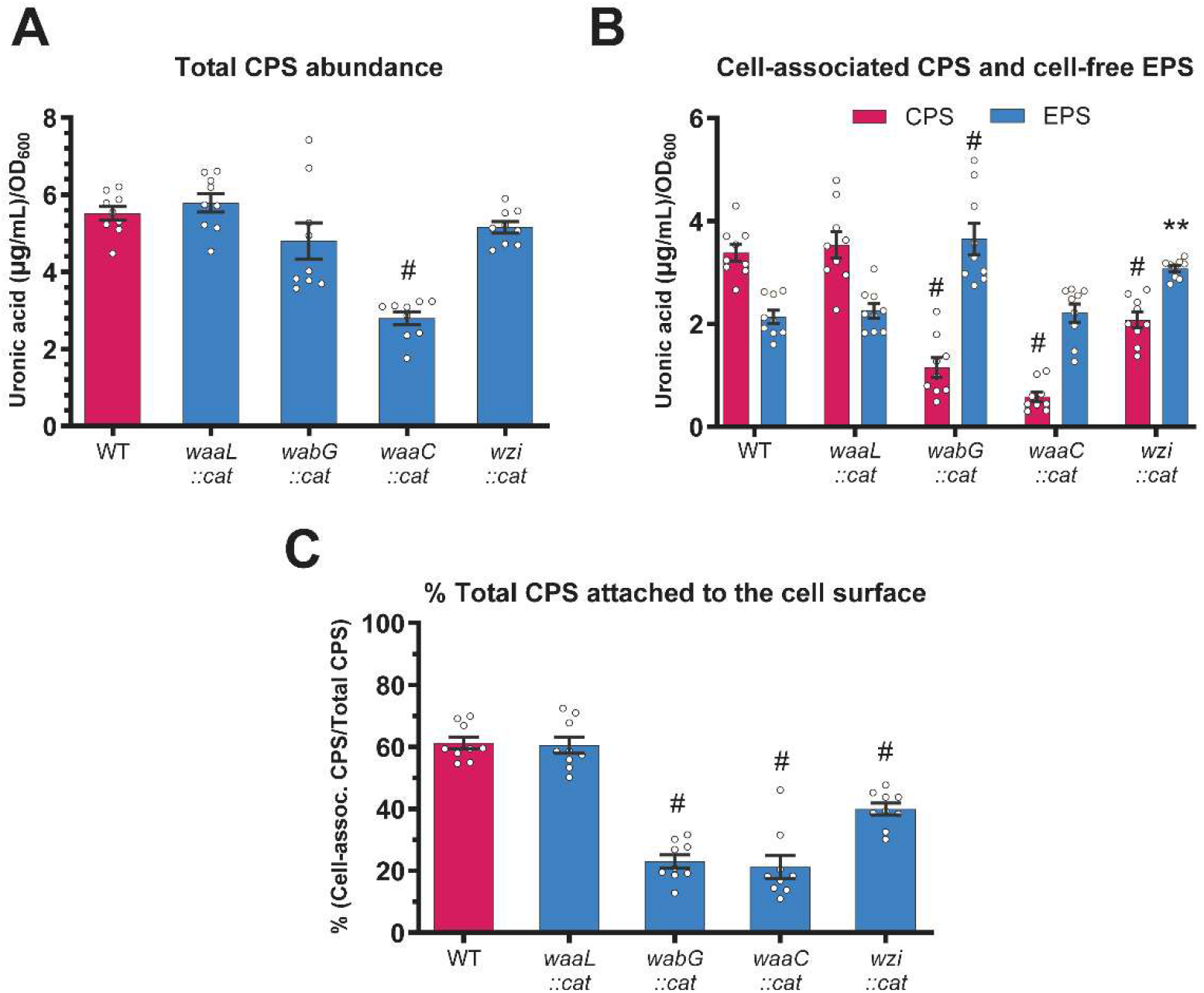
A classical *K. pneumoniae* strain, MKP103, requires LPS-core and Wzi for complete CPS retention. WT and transposon mutants (*waaL::cat*, *wabG::cat*, *waaC::cat* and *wzi::cat*) of *K. pneumoniae* strain MKP103 were cultured in LB and their cell-associated CPS and cell-free EPS abundance and growth kinetics were analyzed. **(A)** Total CPS abundance was determined by measuring uronic acid content in total crude CPS extracts and normalized to OD_600._ **(B)** Cell-associated CPS and cell-free EPS abundance were determined by uronic acid quantification. Cell-associated CPS was calculated by subtracting cell-free EPS from total CPS abundance. **(C)** The percentage of total CPS retained on the bacterial cell surface was calculated as the ratio of cell-associated CPS to total CPS shown in (B). Data presented are the mean, and error bars represent the standard error of the mean. Statistical significance was determined by comparing each mutant to WT using (A,C) one-way ANOVA with Dunnett’s post-hoc test and (B) two-way ANOVA with Dunnett’s post-hoc test. *P*-values are displayed above the bars; ** indicates *p* ≤ 0.01, # indicates *p* ≤ 0.0001. Experiments were performed three independent times, each in biological triplicate (*n* = 9).

**S4 Fig:**
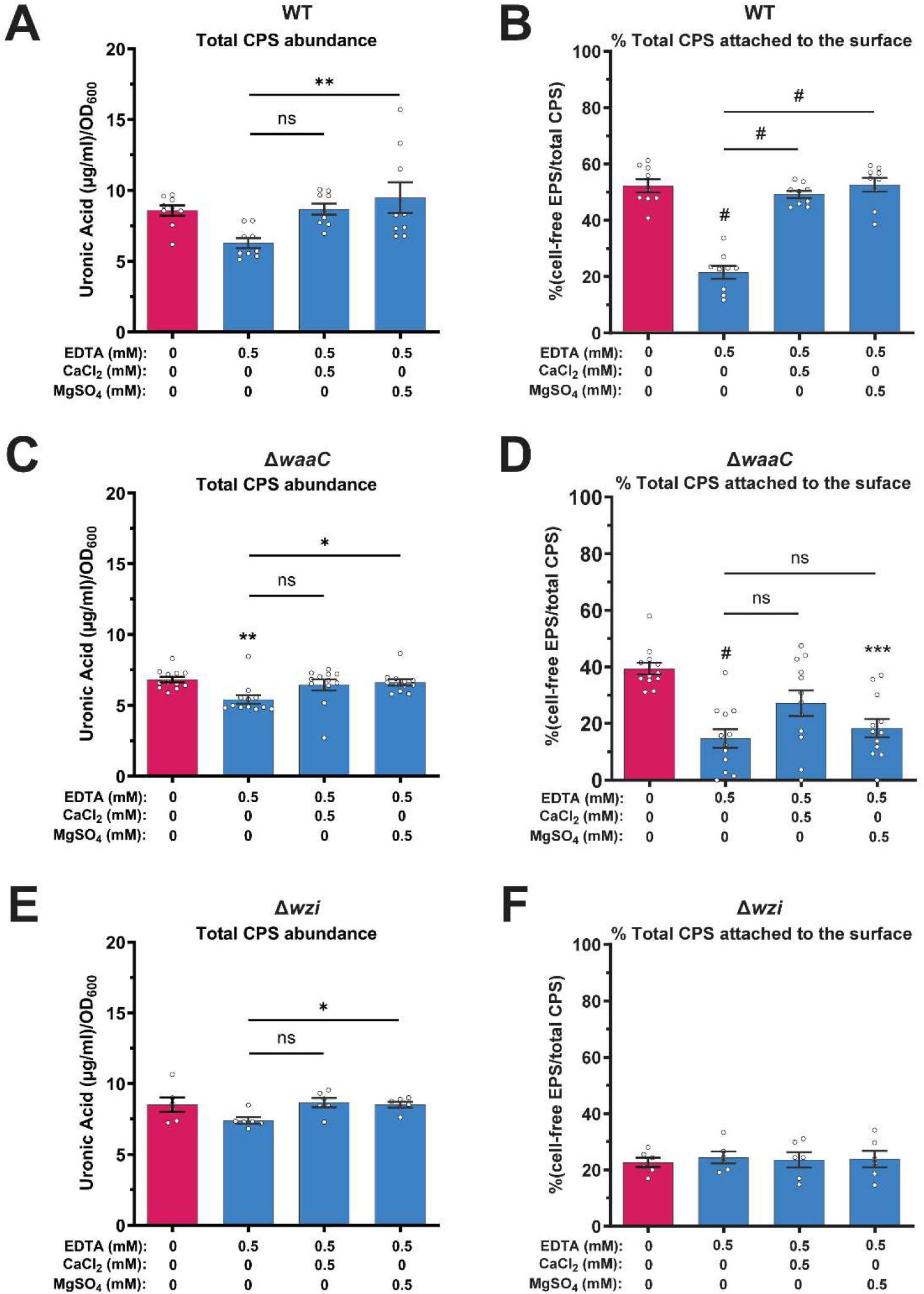
Ca^2+^ and Mg^2+^ supplementation restores EDTA-induced loss of CPS retention in WT, but not in Δ*waaC* or Δ*wzi*. KPPR1 **(A–B)** WT, **(C–D)** Δ*waaC* and **(E–F)** Δ*wzi* were cultured in LB treated with EDTA alone or EDTA supplemented with CaCl_2 o_r MgSO_4._ **(A,C,E)** Total CPS abundance was determined by measuring uronic acid content in total crude CPS extracts and normalized to OD_600._ **(B,D,F)** The percentage of total CPS retained on the bacterial cell surface was calculated as the ratio of cell-associated CPS to total CPS shown in (A,C,E). Data presented are the mean, and error bars represent the standard error of the mean. Statistical significance was determined by two-way ANOVA with Šídák multiple-comparisons test, comparing (i) EDTA-only and EDTA+cation conditions to the untreated condition (*p*-values shown above each bar) and (ii) EDTA+cation conditions to the EDTA-only condition (*p*-values shown by connected lines); * *p* ≤ 0.05, ** *p* ≤ 0.01, *** *p* ≤ 0.001, # *p* ≤ 0.0001. Experiments were performed three independent times, each in biological triplicate (*n* = 9).

**S5 Fig:**
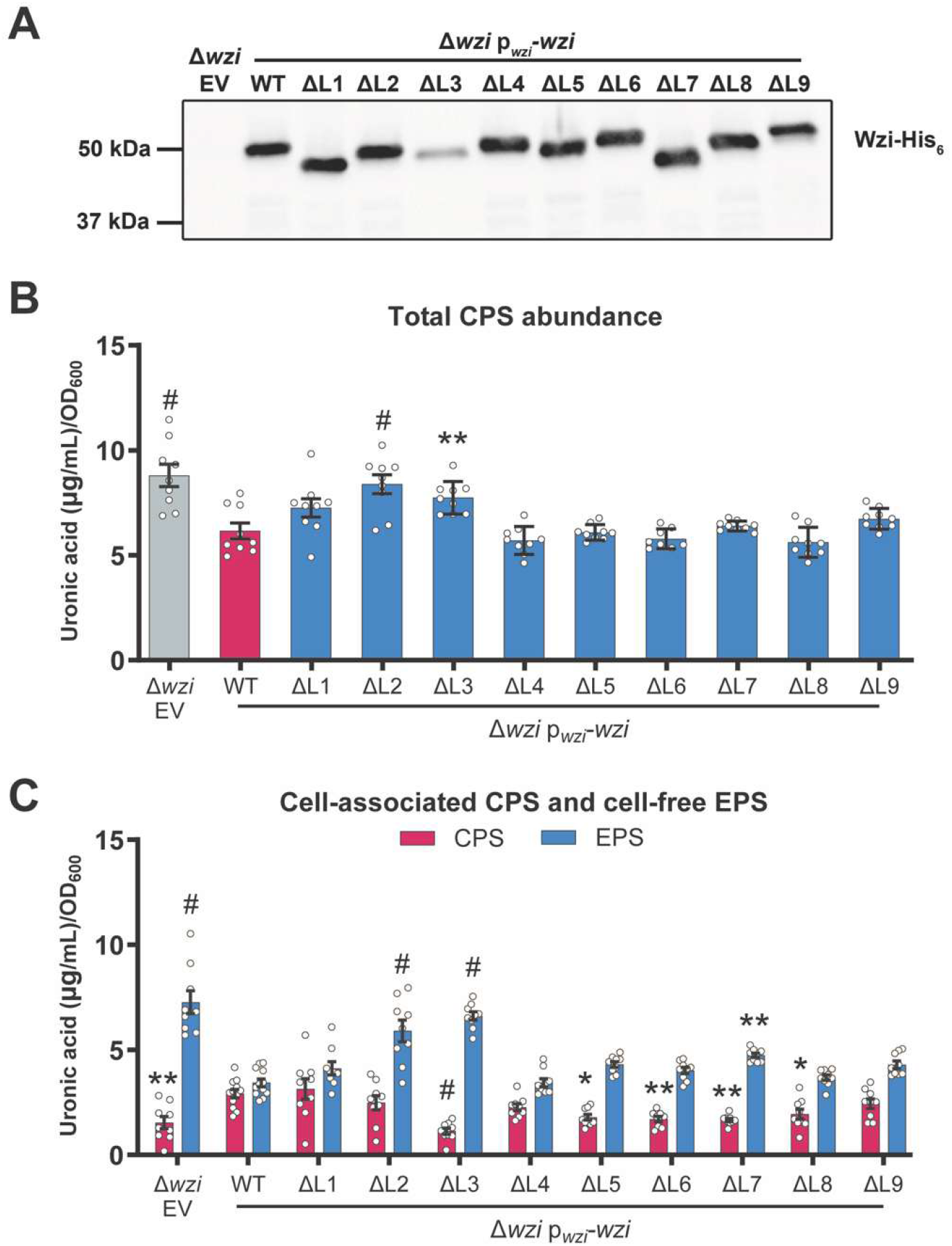
Cell-associated CPS and cell-free EPS of various Wzi loop substitution variants. KPPR1 Δ*wzi* with pBBR1-MCS5empty vector (EV) or the vector expressing WT or loop-substituted *wzi* (p*_wzi_*_-_*wzi*) were incubated in LB. Wzi protein expression was examined and measured for total CPS abundance and cell-free EPS abundance. **(A)** Total cell lysate was prepared from an overnight culture of each strain, resolved by SDS-PAGE and subjected to Western blot, probing for his-tagged Wzi using mouse anti-his HRP-conjugated antibody. **(B)** Total CPS abundance was determined by measuring uronic acid content in total crude CPS extracts and normalized to OD_600._ **(C)** Cell-free EPS abundance was determined by measuring uronic acid content is cell-free supernatant, whereas cell-associated CPS was calculated by subtracting cell-associated EPS from total CPS abundance presented in (A). The corresponding ‘% total CPS attached to the cell surface’ is presented in Fig 3F. Data presented are the mean, and error bars represent the standard error of the mean. Statistical significance was determined by comparing strains expressing loop-substituted *wzi* or EV to the strain expressing WT *wzi*, using (A) one-way ANOVA with Dunnett’s post-hoc test or (C) two-way ANOVA with Dunnett’s post-hoc test. *P*-values are displayed above each bar; * indicates *p* ≤ 0.05, ** indicates *p* ≤ 0.01, # indicates *p* ≤ 0.0001. Experiments were performed three independent times, each with biological triplicate (*n = 9*).

**S6 Fig:**
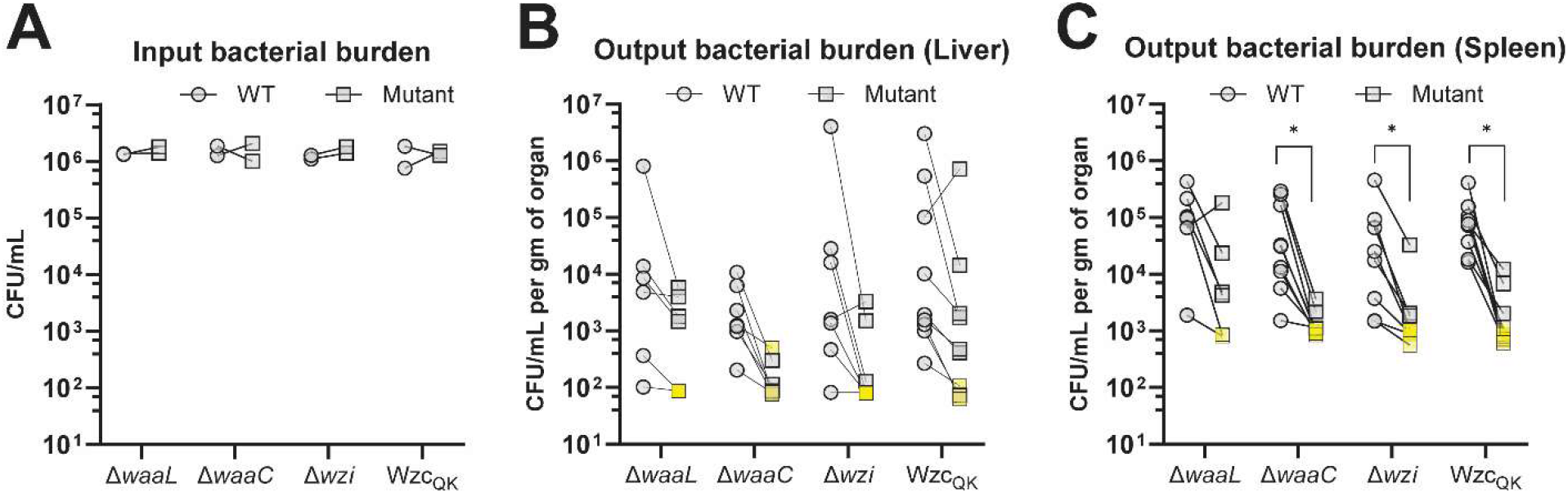
*K. pneumoniae* CPS attachment mutants exhibit reduced liver and splenic burden following transient bacteremia. **(A)** C57BL/6 mice were infected by tail vein injection with a mixture of KPPR1 WT and mutant at 1:1 ratio (10^5^ CFU/mL each). **(B–C)** After 24 hours, bacterial burden in the liver and spleen were enumerated by differential plating. The data presented here are the absolute CFU/mL counts used to calculate log_10 c_ompetitive index in Fig 5 using Equation 2. Each dot represents the CFUs of WT or a mutant. The paired input or output CFUs enumerated from the same mouse are connected by a line. CFUs below the detection limit are indicated by crossed symbol colored in yellow. **(B–C)** Statistical significance was determined using multiple Wilcoxon test with Holm-Šídák correction and calculated by comparing CFU of mutant to WT from the respective group. *P*-values are displayed above each comparison. * *p* ≤ 0.05.

